# Media Selection and Fed-Batch Fermentation Strategy for dsRNA Plasmid Production in *Escherichia coli* GT115

**DOI:** 10.1101/2023.03.07.531461

**Authors:** Adi Pancoro, Intan Taufik, Sena Wijayana

**Author notes:** Corresponding Author Dr. Adi Pancoro & Dr. Intan Taufik, School of Life Sciences and Technology, Bandung Institute of Technology, Ganesa Street No.10, Lb. Siliwangi, Coblong, Bandung, West Java, Indonesia 40132, E-mail address. This research offers novelty about the optimum media for production of pCDNA (+) plasmid containing dsRNA gene in *Escherichia coli* GT115 and its large-scale optimization. Production of plasmid resulted in high plasmid yield. I included it in industrial microbiology’s scope about fermentation and production of synthetic biology. We choose Journal of Industrial Microbiology and Biotechnology (JIMB) because of its impact factor to make this research useful to others.

## Abstract

Application of plasmid encoding synthetic dsRNA targeted IMNV genome (Infectious Myonecrosis Virus) can reduce viral replication in the shrimp industry by activating RNA interference (RNAi) response. Application of dsRNA plasmid as antiviral for IMNV in shrimp-farm need a huge quantity of plasmid. Bioreactor can be used for large-scale plasmid production to achieve high plasmid yield. Plasmid production in the bioreactor can be improved by selection of the host organism, the recombinant plasmid vector, the fermentation media, and the fermentation strategy. This research aim is to determine the fermentation media and fermentation strategy to produce recombinant dsRNA plasmid with high plasmid yield. Selection of fermentation media was conducted in a baffled flask with three different media. Then, the optimum media was used for optimization in bioreactor production with the addition of feeding media. As a result, plasmid production in TB media has a higher biomass growth rate and plasmid production rate than production in M9+Mod and LB+ media. Plasmid production in TB media in baffled-flask resulted in plasmid yield in 2.318 mg/L, 14-fold higher than M9+Mod (0.165 mg/L), and 34-fold higher than LB (0.068 mg/L). In bioreactor production, plasmid production in fed-batch fermentation in bioreactor resulted plasmid yield in 1.018 mg/L, 5-fold higher than batch fermentation (1.882 mg/L). Plasmid was confirmed in agarose gel electrophoresis at ∼5750 bp and insert gene at 700 bp. The cultivation technique developed should be workable for the pilot scale. Downstream processing in plasmid production should be able to achieve plasmid with high concentration and purity.

## Introduction

Necrosis in *Litopenaeus vannamei* tissue is caused by Infectious Myonecrosis Virus (IMNV). IMNV infection drive an annual loss in the Indonesian shrimp industry by 95.6 million USD during 2006-16 (Bondad-Reantaso & Arthur, 2018; Lightner et al., 2012). IMNV is a virus with an 8.2 kbps double-stranded RNA (dsRNA) genome comprising of ORF1 and ORF2 (Naim et al., 2015). Viral infection outbreak within the industry comes up because of the absence of adaptive and specific immune responses. The previous study shows that the transcription of IMNV mRNA can be reduced by RNA interference (RNAi) response, which recognizes the dsRNA fragment encoding IMNV (Rajendran et al., 2022). The development of dsRNA that comprise of the IMNV sequence as an antiviral agent against IMNV infection has emerged recently. A study conducted by Loy, et al. (2013) reveals that the reduction of IMNV copy-number within shrimp body can be achieved after introducing dsRNA fragments coding IMNV gene (Loy et al., 2012; 2013). Therefore, a recombinant dsRNA plasmid encoding the IMNV sequences can serve as a therapeutic and preventive agent against IMNV infection. Other than that, the application of recombinant plasmid also increases dsRNA stability during the introduction process into the shrimp (Itsathitphaisarn et al., 2016).

Plasmid as antiviral cannot be applied with injection into individual shrimp due to time-consuming, labor intensive and impractical for farm-scale application. Delivery of plasmid in shrimp-farm can be applied by oral administration. In addition, use of plasmid as antiviral depends on the stability and availability of plasmid during and after delivery. It can be solved by development of dsRNA plasmid production pipeline and oral administration method (Itsathitphaisarn et al., 2016). Manufacturing recombinant plasmid begins on a laboratory scale by the construction and selection of vector plasmid and bacterial host, followed by optimization of fermentation condition (upstream processing), production and harvesting, then purification steps (downstream processing) (Ferreira et al., 2000). Optimization of plasmid production in the bioreactor can be done by controlling the physical and chemical parameters (pH, dissolved oxygen, and temperature) during the fermentation process, as well as selecting fermentation media and fermentation strategy (Krause et al., 2010).

Culture media selection is important for industrial-scale plasmid production because media is the main component that contributes to production cost. The culture media selection can affect the performance of the microbial process, including plasmid replication. Complex media can increase cell density, but there is a risk of contamination. On the other hands, defined media usually require more components, difficult to prepare and lead to lower cell density than complex media. Culture media selection must be done by considering the plasmid production yield (Danquah & Forde, 2007). Culture media that known to have high-yield plasmid production during the fermentation process are Terrific broth (TB) and minimal salt media, M9 (Galindo et al., 2016; Trivedi et al., 2014). TB is a complex media that can produce plasmid ten-fold higher than the common media for *E. coli* recombinant, Luria-Bertani (LB). The plasmid yield in TB media was achieved in 4.3 mg/L and 11.4 mg/L (Danquah & Forde, 2007; Galindo et al., 2016). Then, M9 is a defined media that can produce plasmid two-fold higher than LB. The plasmid yield in M9 media was achieved in (Trivedi et al., 2014). Silva et al. (2012) also state that the addition of feeding media in fed-batch fermentation can increase plasmid yield during the fermentation process and resulted at two-fold higher than batch fermentation (Silva et al., 2012b). Hence, the objective of this study is to determine the optimum fermentation media between Luria-Bertani, Terrific-broth and M9 minimal media and fermentation strategy with nutrition addition to produce a high yield of dsRNA plasmid in baffled-flask and bioreactor.

## 1. Materials and Methods

### Bacterial strain and plasmid

Bacterial strain used in this research was *Escherichia coli* GT115 (InvivoGen, USA) [F-mcrA ∆ (mrr-hsdRMS-mcrBC) φ80lacZ∆ M15 ∆ lacX74 nupG recA1 araD139 Δ(ara-leu)7697 galE15 galK16 rpsL(StrA) endA1 ∆ dcm uidA(∆ MluI)::pir-116 ∆ sbcC-sbcD] with 5750 bp pIMSY plasmid derived from pCDNA3.1(+) by the addition of dsRNA sequence encoding IMNV gene (dsRNA binding domain, putative fiber 1, 2 and MCP). The bacteria strain had no sbcC and sbcD genes which are beneficial to maintain the plasmid in a hairpin-loop formation. This recombinant bacteria was named by *E. coli* pIMSY

### Laboratory-scale media selection

*E. coli* pIMSY were grown in three different media including Luria Bertani (LB+), Terrific broth (TB) (Himedia), and M9+Modification (M9+Mod) media derived from Cai, et al., (2016) (Cai et al., 2016). LB was added with 4 g/L KH_2_PO_4_, 4 g/L KH_2_PO_4_, and 1.2 g/L (NH4)_2_SO_4_ (MERCK). TB media was added with 0.4% (v/v) glycerol (Vivantics). M9+Mod media formulation was from M9+ formula without choline chloride and riboflavin in MEM Vitamin solution. Cultivation was conducted in baffled flask at 37°C and 150 rpm for 10 h, and 100 ppm ampicillin. Cell culture was collected every 2 h to measure the biomass, pH and plasmid concentration. The biomass concentration (CFU/mL) was determined using a spectrophotometer at 600 nm absorbance while plasmid was isolated using Geneaid-Presto™ Mini Plasmid Kit protocol.

### Batch and fed-batch fermentation in bioreactor

Pilot-scale production of pIMSY was conducted in a 10-L bioreactor School of Life Sciences and Technology, Bandung Institute of Technology, Indonesia. Fermentation was performed on a TB media containing 100 ppm ampicillin. For both batch and fed-batch fermentation, the temperature was maintained at 30°C and pH 7.0. The bacterial strain grown at 30°C to reduce metabolic burden during the fermentation (Gonçalves et al., 2012). Dissolved oxygen (DO) was controlled at 30% using 0.5-1.5 m^3^/h airflow and 150-350 rpm agitation. Batch fermentation was run for 10 h, while fed-batch fermentation 12 h with addition of feeding media containing 400 g/L glycerol (Vivantics), 80 g/L tryptone (Himedia), 10 mL MgSO_4_ 1 M and 100 ppm ampicillin. The feeding media was added to the bioreactor tank after 6 h fermentation, with 200 mL/h feeding rate. Glycol-based anti-foam was added to control foam formation. Culture media was collected every 2 hours, as mentioned earlier.

### Plasmid and dsRNA gene confirmation

pIMSY plasmid was used as a template for restriction and Polymerase Chain Reaction (PCR). The restriction was conducted using Thermo Scientific™ FastDigest NdeI protocol. Meanwhile, PCR was conducted using CMV-F primer (5’-CAA AAT GTC GTA ACA ACT CCG-3’) and BGH-R primer (5’-ATT AGG AAA GGA CAG TGG GA-3’). The condition for PCR amplification was as follows; 5 min at 94°C, 35 cycles of 30 s at 94°C, 30 s at 53°C, and 1 min at 72°C and extension at 72°C for 7 min. Restricted plasmid and PCR product fragment were confirmed by gel agarose electrophoresis.

## 2. Results

### Culture media selection

The growth profile, pH change and plasmid concentration of 150 mL *E. coli* pIMSY culture in baffled-flask with TB, LB+ and M9+Mod media are shown in Figure 1a-c. Growth parameter is shown in Table 1. All cultures were started at the same biomass concentration (10^7^ CFU/mL) and ended at 1.7–2.7 × 10^8^ CFU/mL. This result demonstrates that *E. coli* pIMSY had a slow growth profile in M9+Mod than LB+ and the fast growth profile, TB (Table 1). The growth of *E. coli* pIMSY in M9+Mod media is still active until 10 h cultivation. This biomass growth profile is supported by pH change and plasmid concentration (Figure 1). The pH in M9+Mod media shows more stable than TB and LB+ with a gentle decrease until 10 h. All cultures had a lowest pH at 6, 1 pH different from the started pH (7). Meanwhile, plasmid concentration at the end of cultivation (10 h) results high in TB, 3.863 ± 0.247 mg/L. Plasmid concentration in M9+Mod is still increasing at 10 h cultivation when the increase of plasmid concentration in TB stopped at 4 h, but the concentration is still lower than TB (Figure 1c).

**Table 1.** Biomass growth and plasmid production parameter

| Scale | Type | Temp. (°C) | Media | Log phase (h) | Specific growth rate ($\text{h}^{-1}$) | Generation time (min) | Plasmid production rate ($\text{h}^{-1}$) | Plasmid yield (mg/L) |
| --- | --- | --- | --- | --- | --- | --- | --- | --- |
| Flask | - | 37 | LB+ | 0 – 6 | $0.4070 \pm 0.002$ | $96 \pm 0.263$ | $0.0698 \pm 0.057$ | $0.068 \pm 0.057$ |
| | - | 37 | M9+Mod | 0 – 10 | $0.3400 \pm 0.008$ | $106 \pm 1.357$ | $0.1479 \pm 0.026$ | $0.165 \pm 0.051$ |
| | - | 37 | TB | 0 – 4 | $0.6604 \pm 0.004$ | $66 \pm 0.241$ | $0.7065 \pm 0.159$ | $2.318 \pm 0.651$ |
| | - | 30 | TB | 0 – 6 | $0.522 \pm 0.005$ | $80 \pm 0.567$ | $0.3465 \pm 0.043$ | $1.808 \pm 0.281$ |
| Bio-reactor | Batch | 30 | TB | 0 – 6 | $0.508 \pm 0.052$ | $81 \pm 6.018$ | $0.3498 \pm 0.022$ | $1.882 \pm 0.784$ |
| | Fed-batch | 30 | TB + feed media | 0 – 12 | $0.422 \pm 0.022$ | $98 \pm 3.152$ | $0.3976 \pm 0.013$ | $10.186 \pm 2.315$ |

Experimental error shows in standard error from duplicated sample.

**Figure 1.**
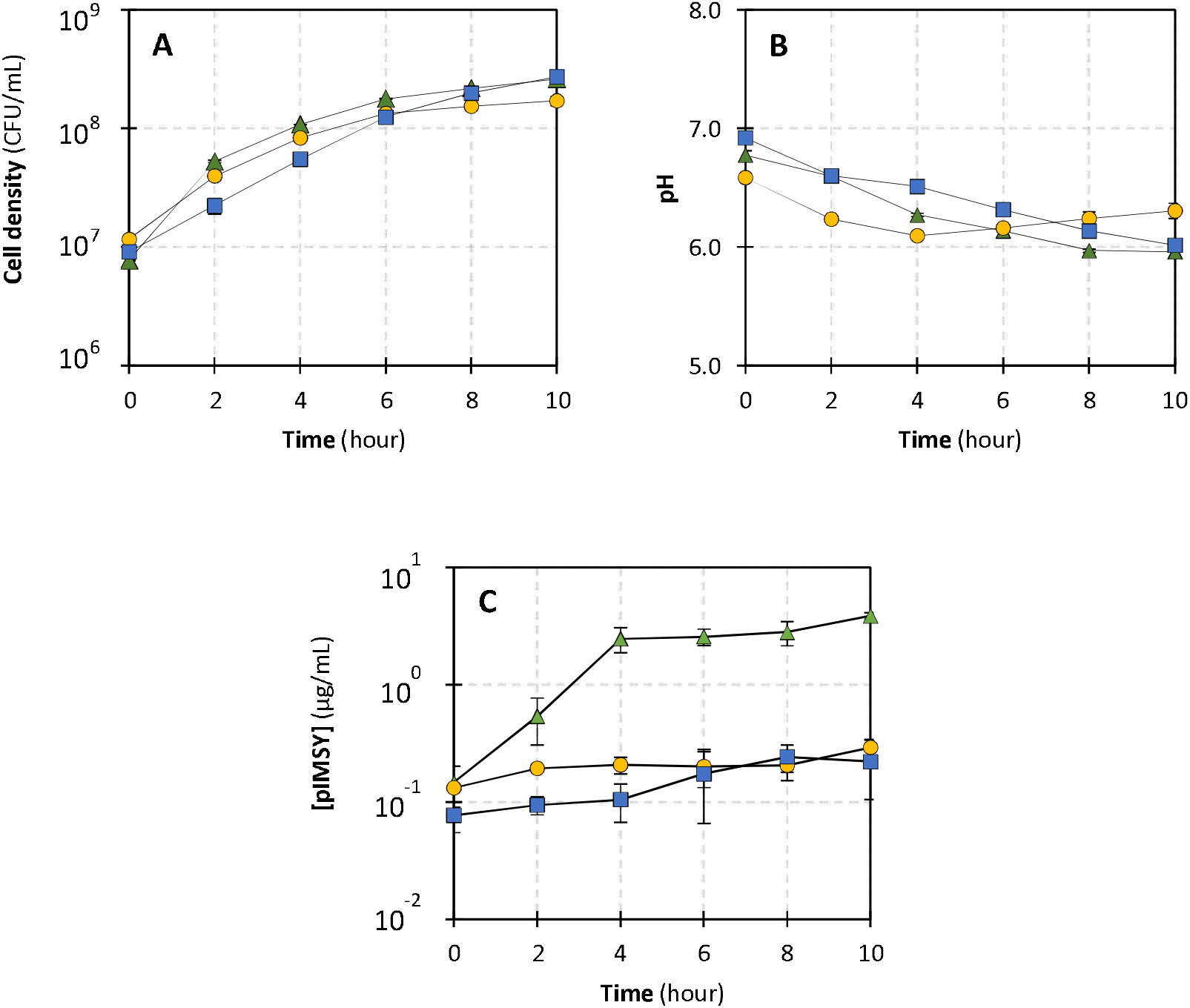
Growth profile of *E. coli* pIMSY in different media. Biomass growth curve (**A**), pH change (**B**) and plasmid concentration (**C**). Cultivation was conducted in LB+ (*yellow-circle*), TB (*green-triangle*) and M9+Mod (*blue-square*) with 100 ppm ampicillin at 37°C and 150 rpm agitation. All cultures were conducted in duplicate. *Error bars* show the experimental error between duplicates.

The growth rate of *E. coli* pIMSY and plasmid production rate is high in TB media. Plasmid production rate in TB is 10-fold higher than LB+ and 5-fold higher than M9+Mod and resulted in high plasmid yield at the end of cultivation. The plasmid yield in TB media cultivation result high in 2.318 ± 0.651 mg/L, 34-fold higher than plasmid yield in LB+ media and 14-fold higher than M9+Mod media (Table 1).

### Plasmid confirmation

The plasmid and dsRNA gene fragment confirmed using gel electrophoresis that was found within ∼5750 bp and 700 bp DNA bands, respectively (Figure 2). There is also another DNA band size was found in isolated plasmid before restriction and PCR product fragment.

**Figure 2.**
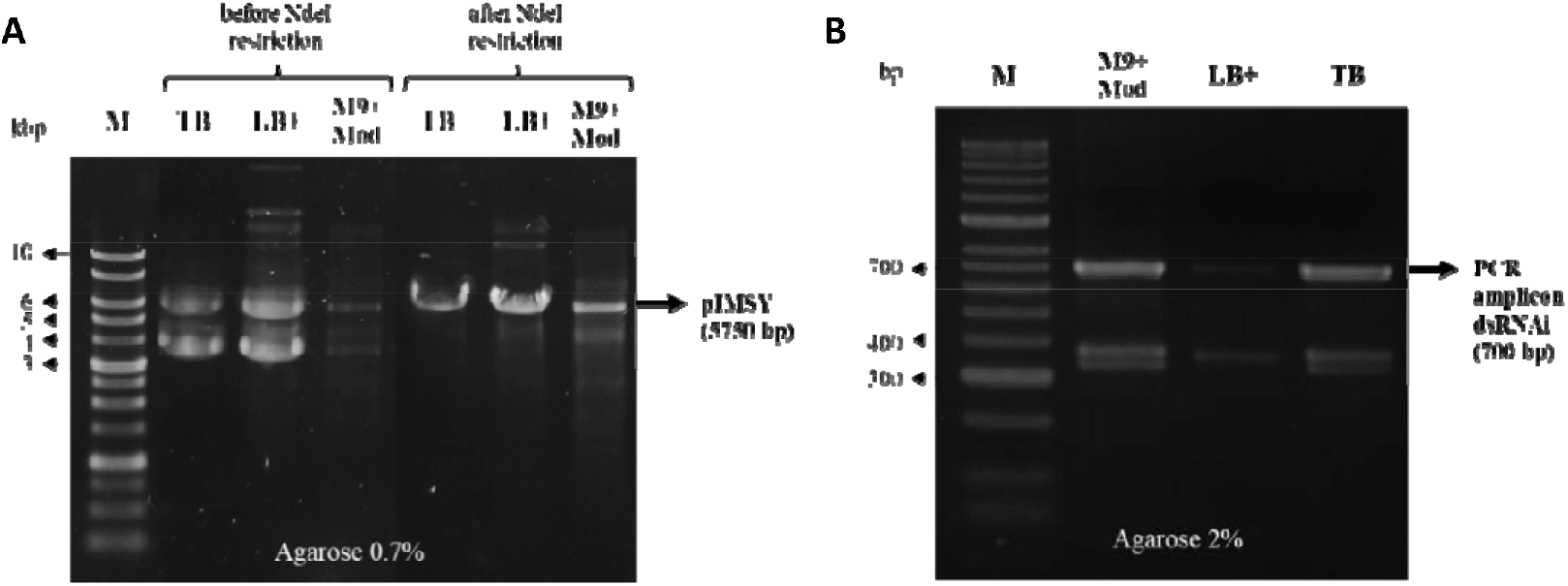
Plasmid and dsRNA gene fragment confirmation in different cultivation media. Agarose gel electrophoresis of isolated plasmid before and after NdeI restriction with 0.7% agarose (**A**) and PCR fragment product of dsRNA gene with 2% agarose (**B**). The isolated plasmid is from 10 h cultivation. M9+Mod = cultivation in M9+Modification media, LB+ = cultivation in Luria-Bertani media, TB = cultivation in Terrific-broth media, M in isolated plasmid electrophoresis was 1 KB DNA Ladder–SMOBIO, then M in PCR fragment electrophoresis was 50 bp DNA Ladder–Bioline. This electrophoresis was conducted in 100 volts and 26 min. Staining method using ethidium bromide (EtBr) and visualized in UV-transilluminator.

### Fermentation in bioreactor with batch and fed-batch fermentation

The growth profile and plasmid concentration of 7 L *E. coli* pIMSY culture in 10 L bioreactor with TB media and different fermentation strategy (batch and fed-batch) are shown in Figure 3a and b. Growth parameter is also shown in Table 1. All cultures were started at the same biomass concentration (10^7^ CFU/mL) but ended at different biomass concentration, 4.2 × 10^8^ ± 1.3 × 10^8^ CFU/mL in batch fermentation and 1.6 × 10^9^ ± 8.2 × 10^7^ CFU/mL. *E. coli* pIMSY resulted in 4-fold higher than biomass concentration with fed-batch fermentation than batch. The growth of *E. coli* pIMSY and plasmid production in fed-batch is still active after feeding media addition after 6 hours of cultivation (until 12 h), when *E. coli* pIMSY in batch fermentation enter the stationary growth phase after 6 h cultivation (Figure 3). Plasmid concentration at the end of cultivation in fed-batch fermentation (12 h) results with 10.273 ± 2.348 mg/L, also four-fold higher than batch fermentation 2.437 ± 0.615 mg/L (Figure 3b).

**Figure 3.**
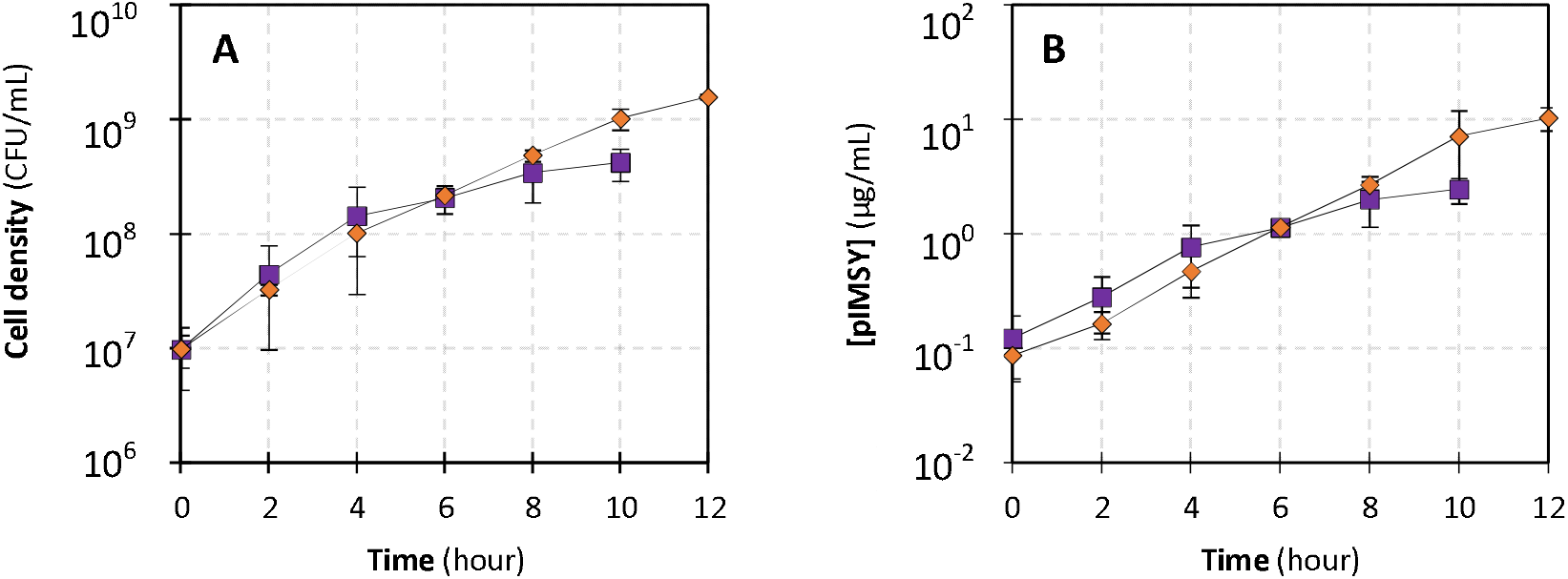
Growth profile of *E. coli* pIMSY in different fermentation strategy. Biomass growth curve (**A**) and plasmid concentration (**B**). Cultivation was conducted in TB (batch) and feeding media (fed-batch) with 100 ppm ampicillin at 30°C and 150-350 rpm agitation. *Orange-diamond* showed in fed-batch fermentation when the *purple-square* showed in batch fermentation. All cultures were conducted in duplicate. *Error bars* show the experimental error between duplicates.

The growth rate of *E. coli* pIMSY and plasmid production rate in batch and fed-batch fermentation are similar. Meanwhile, because the plasmid production in batch fermentation stopped at 6 h of cultivation and still active in fed-batch fermentation until 12 h of cultivation. The plasmid yield in fed-batch fermentation result high in 10.186 ± 2.315 mg/L, 5-fold higher than batch fermentation (Table 1).

## 3. Discussion

### TB resulted in high plasmid yield

Medium selection is an upstream of plasmid production process to determine the cultivation media with resulted in high plasmid yield. The parameter for culture media selection based on bacterial growth and plasmid production performance. The bacterial growth parameters are the period of log phase, specific growth rate, generation time and biomass yield. Then, the plasmid production parameters are plasmid production rate and plasmid yield (Berenjian, 2019).

*E. coli* pIMSY growth rate resulted high in TB media cultivation, followed by LB+ and M9+Mod (Table 1). Terrific broth is a complex medium of *E. coli* cultivation contains high content of complex nitrogen and carbon sources. Compared to LB+, TB contains yeast extract and tryptone concentration with 20 and 480% higher. TB also contains phosphate buffer that can maintain pH still stable and give high buffering capacity and glycerol as an extra and defined carbon source. The high nutrition content in TB makes *E. coli* easier to get substrate for growth and leads *E. coli* growth faster in TB (Galindo et al., 2016). The complex nutrition in TB and LB+ from yeast extract and tryptone comprises various types and forms of carbohydrates, nucleic acids, proteins, minerals, and vitamins that can be uptake by *E. coli* and involved in every metabolism pathway. It makes *E. coli* easier to get substrate for growth with molecule transport mechanism than synthesis all of them (Ogoshi et al., 2014).

*E. coli* pIMSY growth rate in M9+Mod was lowest because of the low concentration of single carbon source (2% (m/v) of glucose) and the nitrogen source was in organic nitrogen (ammonia). *E. coli* can uptake glucose with phosphonyl pyruvate (PEP) phosphotransferase system (PTS) like ATP-binding cassette (ABC) transporter and also porin protein transporter LamB (Natarajan & Srienc, 1999). The glucose molecules will be used as an energy source with glycolysis, pyruvate decarboxylase, tricarboxylic acid (TCA) cycle and oxidative phosphorylation pathway in aerobic condition and resulted adenosine triphosphate (ATP), acid, and ethanol during oxygen limitation (Han et al., 1992). *E. coli* also can uptake ammonia in M9+Mod with transporter protein AmtB as nitrogen source in aerobic condition to produce glutamate and glutamine by glutamate dehydrogenase- (GDH) or glutamine synthetase- (GS) dependent pathway (Reitzer, 2003). This in organic nutrition cannot be used directly as *E. coli* growth substrate and it must be processed in organic nitrogen synthesis like amino acid and nucleic acid. *E. coli* also can synthesize an amino acid or nucleic acid form derivate molecules in glucose metabolism such as acetyl co-A as glutamine precursor (Ginésy et al., 2017). This process also needs more energy than using the uptake of organic nitrogen source in culture media. Metabolism process in synthetic media lead the growth of *E. coli* in M9+Mod slower than complex media (LB+ and TB). The ends of cell density in synthetic can be achieved like in complex media with longer time cultivation (Berenjian, 2019).

Vitamin also important nutrition for bacterial growth. *E. coli* can uptake extracellular vitamin by transporter protein like ATP-binding cassette (ABC) transporter, abgT transporter, and energy-coupling factor (ECF) transporter (Carter et al., 2007; Du et al., 2011; Vallari & Rock, 1985). M9+Mod also added by vitamin such as pantothenate (B_5_ vitamin) and myo-inositol for acetyl co-A precursor in energy metabolism before TCA cycle, folic acid (B_9_ vitamin) for a precursor in folate metabolism including nucleic acid and amino acid biosynthesis, pyridoxine (B_6_ vitamin) for coenzyme amino acid, glucose, and lipid metabolism, thiamine (B_1_ vitamin) for coenzyme in sugar and amino acid catabolism, then nicotinic acid for NAD and NADP precursor (Nancib et al., 1991). These vitamins also contain in yeast extract with different concentration (Ogoshi et al., 2014). Based on the result, vitamin composition and concentration in M9+Mod (about 1-2 ppm) does not result in high yield plasmid. It can be hypnotized that composition and concentration in the TB media are favorable for plasmid production.

All culture media resulted in decrease pH because of acid compound as metabolic product In *E. coli*, such as acetic acid and lactic acid. Acetic acid production is known in high at complex media (LB+ and TB) than defined media (M9+Mod) that lead a significant of pH decreases (Losen et al., 2004). There is also pH increasing after 4 h cultivation in LB+ (Figure 1) because of acetic acid consumption in carbon source limitation. LB+ contain carbon source lower than TB that can lead to carbon limitation. Here, *E. coli* used acetic acid as a carbon source with glycosylation pathway (anabolism) to changes acetic acid into acetyl co-A (Liria et al., 1998). Meanwhile, there is a higher yeast extract concentration in TB and added by glycerol that can make TB have a lot of carbon source than LB+. The pH changes in M9+Mod seems more stable because of high buffer capacity from buffer solution (KH_2_PO_4_, K_2_HPO_4_ and Na_2_HPO_4_) (Cai et al., 2016).

Plasmid production in all culture media resulted in the same trend as biomass growth. It means that plasmid was a primary metabolic in *E. coli*. Plasmid as extrachromosomal DNA in *E. coli* pIMSY gives more metabolic burden during plasmid replication and AmpR gene expression as a selectable marker. This metabolic burden lead *E. coli* needed more energy source to growth. It makes *E. coli* pIMSY resulted high plasmid yield in TB than M9+Mod and LB+ (Table 1). TB has more energy source and contains amino acid and vitamin from tryptone also phosphorous ion that is important for a precursor in nucleic acid biosynthesis (Galindo et al., 2016; Ogoshi et al., 2014).

### Plasmid and dsRNA gene fragment confirmation

Based on the gel electrophoresis, there are three DNA bands in isolated plasmid from all cultivation media before NdeI restriction because of the different conformation of plasmid (supercoil, circular and linear) (Figure 2a). It was confirmed by the result after NdeI restriction with one DNA band in gel electrophoresis (Figure 2a) at plasmid size (5750 bp). NdeI restriction makes all conformation of isolated plasmid becoming a linear conformation (Tirabassi, 2021). Then, the gel electrophoresis from the PCR fragment resulted in two DNA bands at 700 and 350 bp. This result happened because of the secondary structure in the PCR fragment because of the inverted repeat in dsRNA gene sequence (Primrose & Twyman, 2006).

### Batch and fed-batch fermentation

The growth of *E. coli* pIMSY can be optimized by nutrition addition before the bacteria enter the stationary phase. The growth of *E. coli* pIMSY at batch fermentation in bioreactor resulted in a stationary phase after 6 h cultivation because of nutrition limitation (Figure 3a) (Nancib et al., 1991). This statement can be proven by nutrition addition (fed-batch) at the end of log phase with feeding media containing 400 g/L glycerol, 80 g/L tryptone, 4% (v/v) MgCl_2_ 1M and ampicillin 100 ppm (Silva et al., 2012a). Glycerol was used for carbon source to reduce production cost and also was the extra carbon source in TB (Tartof & Hobbs, 1987; Trchounian & Trchounian, 2015). It reported that feeding media addition can maintain the growth of *E. coli* pIMSY still active until 12 h cultivation (Figure 3a).

At the feeding media addition, *E. coli* used glycerol as a carbon source for response of carbon source limitation from yeast extract. *E. coli* can transport extracellular glycerol by facilitated diffusion mechanism with membrane integral protein GlpF. The metabolism of glycerol is related to glucose, intracellular glycerol will be converted into glycerol-3-phosphate (G3p) by glycerol kinase (GlpK). This G3p molecule can be converted into dihydroxyacetone-phosphate (DHAP) as a precursor of pyruvate. Then, pyruvate can be used for energy source for cell division with conversion into acetyl co-A, acetic acid, or lactic acid (Murarka et al., 2008).

## Conclusion

This report described the method of using a cultivation approach for pilot-scale production of plasmid containing dsRNA sequence targeting multiple IMNV genes. First, cultivation in the Terrific broth medium resulted in higher plasmid yield than M9+Modification and Luria Bertani+ medium. The amount of plasmid as determined from the Terrific broth medium was 34-fold higher than the yield obtained under Luria Bertani+ medium and 14-fold higher than M9+Modification medium. For pilot-scale production, fed-batch fermentation with glycerol, tryptone and MgCl_2_ should be suitable for the pilot-scale of plasmid dsRNA production. The amount of plasmid as determined from the fed-batch culture was 5-fold higher than the yield obtained under the batch fermentation.

Extend the addition time of feeding medium during fed-batch fermentation should be able to maintain biomass growth and plasmid production in the logarithmic phase and result in high plasmid yield production.

## Credit author statement

**Dr. Adi Pancoro & Dr. Intan Taufik:** Supervision, Methodology, Conceptualization, Resources, Funding acquisition, Writing – Review & Editing.

**Sena Wijayana:** Conceptualization, Investigation, Methodology, Formal analysis, Validation, Writing – Original Draft.

## 4. Acknowledgements

The authors thank Dr. Neil Priharto (Head of Bioprocess Laboratory, School of Life Science and Technology, Bandung Institute of Technology) for supervision in bioreactor.

